# Parallel encoding of information into visual short-term memory

**DOI:** 10.1101/398990

**Authors:** Edwin S. Dalmaijer, Sanjay G. Manohar, Masud Husain

## Abstract

Humans can temporarily retain information in their highly limited short-term memory. Traditionally, objects are thought to be attentionally selected and committed to short-term memory one-by-one. However, few studies directly test this serial encoding assumption. Here, we demonstrate that information from separate objects can be encoded into short-term memory in parallel. We developed models of serial and parallel encoding that describe probabilities of items being present in short-term memory throughout the encoding process, and tested them in a whole-report design. Empirical data from four experiments in healthy individuals were fitted best by the parallel encoding model, even when items were presented unilaterally (processed within one hemisphere). Our results demonstrate that information from several items can be attentionally selected and consequently encoded into short-term memory simultaneously. This suggests the popular feature integration theory needs to be reformulated to account for parallel encoding, and provides important boundaries for computational models of short-term memory.

## Introduction

Because the world offers more visual information that humans can process, only a subset can be selected (Baddeley & Hitch, 1974; Posner, Snyder, & Davidson, 1980). Attention is often compared to a spotlight that selects information from a small sector of the visual field before shifting to other locations (Crick, 1984; Duncan, 1984; Eriksen & St. James, 1986; Saarinen & Julesz, 1991). According to the popular feature-integration theory, only attended information can be bound together and committed to visual short-term memory (Treisman & Gelade, 1980; Wheeler & Treisman, 2002), where items’ features are stored in hierarchical bundles (Brady, Konkle, & Alvarez, 2011).

Although much research has focussed on the architecture of visual short-term memory storage (Bays, 2015; Bays & Husain, 2008; Luck & Vogel, 1997; Oberauer & Lin, 2017; Wilken & Ma, 2004; Zhang & Luck, 2008), little work has been conducted on the encoding mechanism, the process by which visual information first enters the short-term store. There exist two main theories: The serial hypothesis is central to feature integration theory, and postulates that items are attended to and consequently encoded one-by-one (Treisman & Gelade, 1980; Wheeler & Treisman, 2002), whereas the parallel hypothesis states that all items simultaneously compete for attention and access to short-term memory (Bundesen, 1990). These theories have informed contemporary computational models of short-term memory that either implicitly assume serial encoding (Manohar, Zokaei, Fallon, Vogels, & Husain, 2017), or allow parallel encoding (Matthey, Bays, & Dayan, 2015).

Despite the prominence of feature-integration theory, its serial encoding assumption has received little empirical testing (Liu & Becker, 2013). Encoding can be studied by varying the duration of the stimuli, and modelling demonstrates that both its rate and capacity are limited (Bundesen, 1990). This has been confirmed in more recent studies that were either agnostic about the encoding mechanism (Bays, Gorgoraptis, Wee, Marshall, & Husain, 2011), or assumed serial encoding without formal verification (Vogel, Woodman, & Luck, 2006).

There is some evidence that features of unattended items may still be bound together into objects (Houck & Hoffman, 1986), and that humans can attend to two non-contiguous locations at the same time from psychophysics (Hahn & Kramer, 1998; Kramer & Hahn, 1995) and neuroimaging (McMains & Somers, 2004; Müller, Malinowski, Gruber, & Hillyard, 2003). Although these findings challenge the serial attending and binding hypothesis in feature-integration theory (Treisman & Gelade, 1980), it remains unclear whether they challenge the serial encoding hypothesis (Wheeler & Treisman, 2002).

One crucial difference between serial and parallel encoding is the probability of *only* one item being in short-term memory when two items are presented simultaneously. Specifically, the serial model predicts a period when it is more likely that only one of the two items has been encoded, because encoding of the second item cannot begin until the first item has completed. In contrast, if encoding occurred in parallel, the probability of each item being encoded should rise together, and thus the probability of only one of the two being encoded at any time is lower (**Figure 1**). Note that in a parallel model, it is still possible for only one item to be in memory, because the rates of encoding the two items might vary stochastically. However, this probability will always be lower than for a comparable serial model.

**Figure 1.**
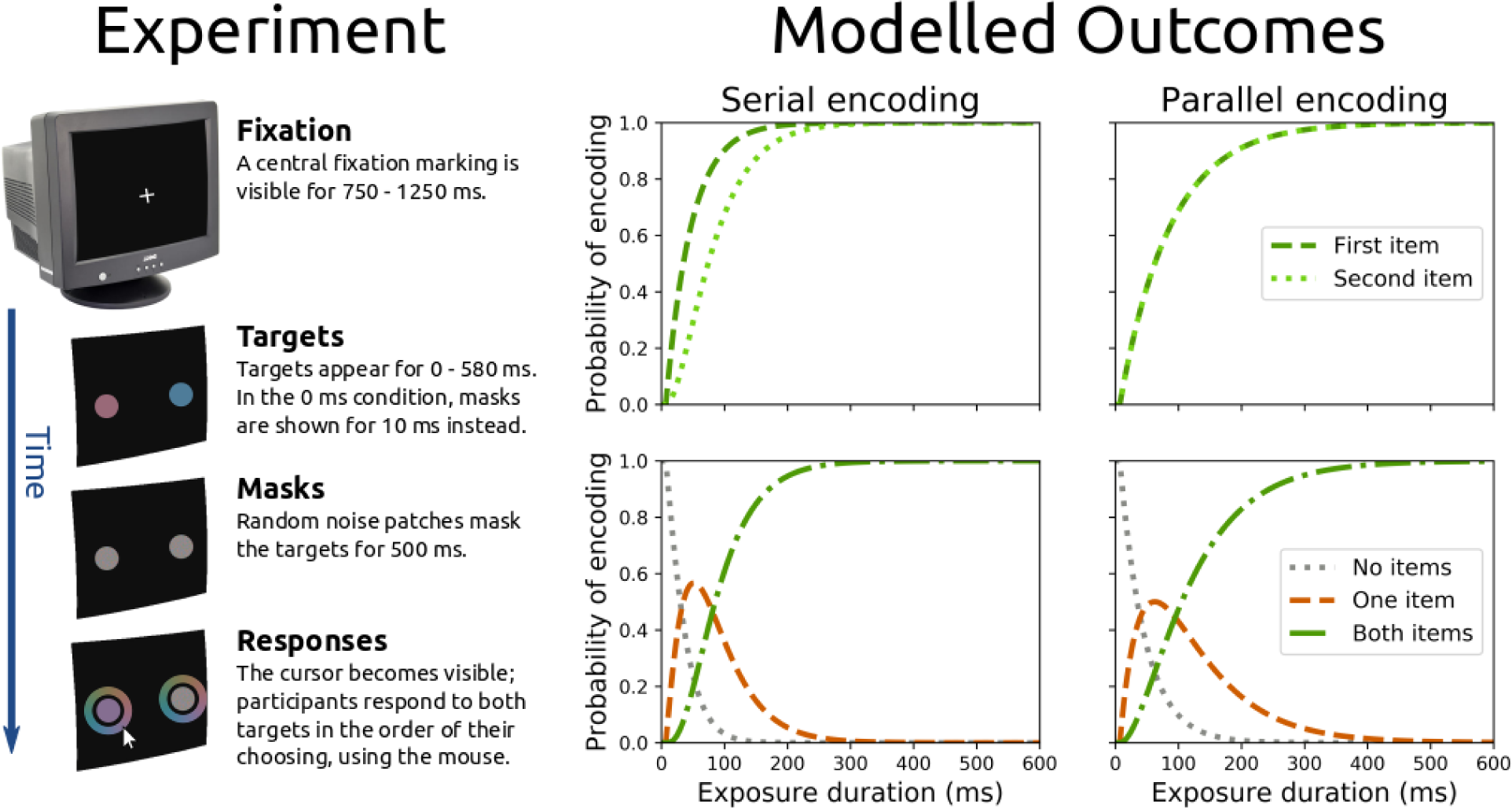
In four experiments, each of the 500-900 trials followed the same pattern of a central fixation marker, followed by one or two targets presented either unilaterally or bilaterally for a variable exposure duration (0-580 ms), after which they were masked by patches of random noise for 500 ms. Participants were required to memorise and respond to both targets. The top row of modelled outcomes represents how the encoding probabilities of each individual item change over time. Crucially, in the serial model the probability of each item being encoded rises sequentially, whereas they rise simultaneously in the parallel model. The bottom row of modelled outcomes shows the combined individual encoding probabilities, and represents the probabilities that no items (grey dotted line), only one item (orange dashed line), or both items (green dashed-dotted line) are encoded. Crucially, it shows that in the serial model, the probability that only one item is encoded peaks more sharply at lower encoding times. (For the modelled outcomes, the encoding rate was set to 25 Hz so that probability of the first item being encoded was 0.76 at 50 ms in a serial scenario, which is in line with empirical estimates (Bundesen, 1990; Vogel et al., 2006). In the serial model, all processing capacity was used to encode one item first, and the other item second. In the parallel model, the processing capacity was divided equally between both items, which were encoded at the same time.)

Here, we describe two competing models for short-term memory encoding: A serial process that encoded one item at a time (**Equation 1-3**), and parallel processes that encodes items at the same time (**Equation 4**). Both models are rate and capacity limited, in line with experimental observations (Bays et al., 2011; Bundesen, 1990; Vogel et al., 2006). Given the probability that each item in a memory array has been encoded by a given time, we can compute the joint probabilities that no items, only one item, and both items have been encoded by that time (**Equations 5-7**).

We denote the time at which the first item is encoded by *T1*, and the second item by *T2*. In the serial model, the probability of the first item being encoded before time *t, P(T*_*1*_*<t),* is given by **Equation 1**, where *κ* is the encoding rate, and *t0* is the minimally effective exposure duration (not unlike (Bundesen, 1990)) that accounts for lag in visual processing. Before the onset of encoding, when *t* is smaller than *t0*, *P* is set to 0.

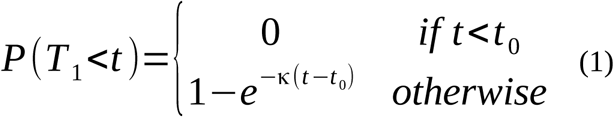

In this serial model, encoding of the second item can only start after encoding of the first item, and thus the probability of the second item being encoded by time *t*, *P(T2<t)*, is determined by the probability that the first item has been encoded, *P(T1<t)*. Processing time for the second item is counted from the encoding of the first, and encoding rate *κ* is the same for each.

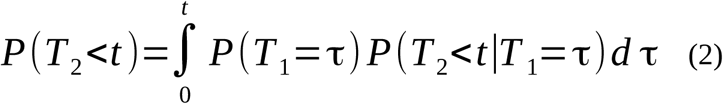

Or:

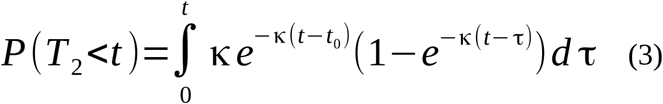

In the parallel model, both the probability of the first item being encoded by time *t*, *P(T*_*1*_*<t)*, and of the second item being encoded, *P(T*_*2*_*<t)*, is given by **Equation 4**.

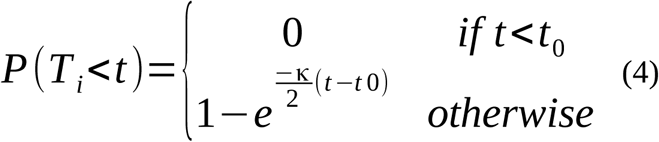

When participants recall several items in the order of their choice, later items are known to be reported less precisely (Adam, Vogel, & Awh, 2017). Therefore, we included a third parameter *α* to account for serial recall, *without* assuming a correspondence between encoding and response order (see Methods).

Both serial and parallel models thus yield three free parameters: rate *κ*, lag *t0*, and second-report asymptote *α*. For each model, the probabilities that both (**Equation 5**), only one (**Equation 6**), or no items are encoded (**Equation 7**) follow directly from the probabilities of either item being encoded:

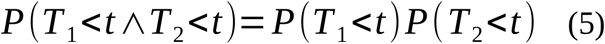

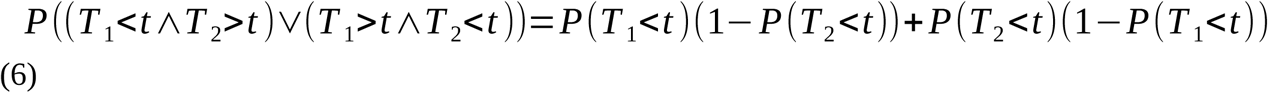

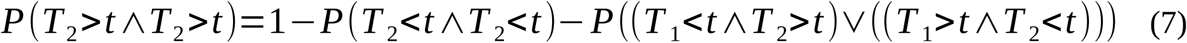

To test the models, we developed a new whole-report paradigm in which we arrest encoding by presenting stimuli for very short durations followed by a mask, but then require participants to report both stimuli. This allowed us to probe whether neither, one, or both items were encoded, after a given encoding time. In each trial, participants were asked to remember up to two items that were presented very briefly (0-580 ms), and to recall the orientation or colour of both items (**Figure 1**). By varying the exposure durations of the items before they were masked, we could probe the probabilities of items being present in short-term memory throughout the encoding process. Here, we present model fits and additional results four experiments in a total of 60 healthy participants.

## Methods

### Procedure

Participants were recruited through the University of Oxford’s Experimental Psychology recruitment website for healthy volunteers, with permission from the local ethics committee. They were compensated at a rate of 8 British Pounds per hour. Each experiment consisted of 500-900 trials, and lasted no longer than two hours. A total of 60 participants took part in four experiments (N=11, N=11, N=16, N=22); numbers were modelled after previous research (Bays et al., 2011; Bundesen, 1990; Vogel et al., 2006).

In all experiments, stimuli were presented on fixed locations for brief durations, and then replaced by masks. In experiments 1, 2, and 4 the targets were coloured disks (sampled from a circle with radius 22 around the whitepoint in CIE L*a*b* space with L=50), and in experiment 3 they were black-to-white sinusoidal gratings (5 cycles per stimulus). In experiment 1, the exposure durations were 0, 10, 20, 30, 50, 90, 140, and 200 milliseconds, and in experiments 2-4 they were 0, 10, 20, 40, 70, 120, 200, 360, and 580 milliseconds. In experiment 1 the stimuli always appeared left and right; in experiments 2 and 3 there could appear one (50% of trials; randomly selected left or right location) or two stimuli (50% of trials; both locations); and in experiment 4 the stimuli could appear unilaterally (top and bottom on the left or right half) or bilaterally (left and right on the top or bottom half). Stimuli were presented at 7.5 degrees of visual angle from the display centre, and had a diameter of 2.7 degrees of visual angle.

Mask duration was 500 milliseconds in all experiments, after which the mouse cursor appeared. At this point, participants could use the cursor to click on a mask (or the surrounding colour wheel), which then reoriented towards the cursor. Participants could fine-tune their response by holding down the mouse button and moving the cursor. After responding to all stimuli in the order of their choice, participants could press the Space key to continue the experiment.

Experiments were programmed in Python (Dalmaijer, 2017; Van Rossum & Drake, 2011), using the PyGaze (Dalmaijer, Mathôt, & Van der Stigchel, 2014) and PsychoPy (Peirce, 2007) packages.

### Data analysis

Error was computed as the circular difference between stimulus colour (angle in colour space) or orientation, and the associated response. The average unsigned error was used as an index of performance per stimulus per exposure duration. Repeated-measures ANOVAs were conducted in JASP, version 0.7 (JASP Team, 2016).

In addition, error distributions for each exposure duration were fitted to mixture models (Bays, Catalao, & Husain, 2009; Zhang & Luck, 2008), using maximum-likelihood estimation. The probability density function of the first response 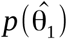 was described by the sum of three components (**Equation 8**). The first component described the responses that were made with the target at the response location in memory, and should fit Von Mises distribution 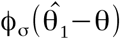 with density parameter *σ* and centre 0 (the distance between response and target orientation under perfect memory). The second component described uniformly distributed random guesses at height 2π. The third component described responses that were made with the target at the other location in mind (“swap error”), and should fit Von Mises distribution 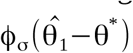. In this framework, the *γ*_*1*_ parameter reflected the proportion of trials in which the first response was a guess, and the *β*_*1*_ parameter reflected the proportion of trials in which the first response was a swap error. The probability density function for the second response (**Equation 9**) took the same form. Each model yielded three free parameters: guessing probability *γ*, swapping probability *β*, and density *σ*.

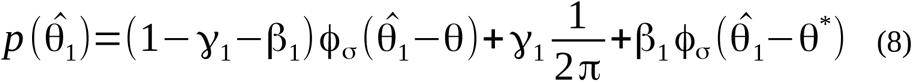

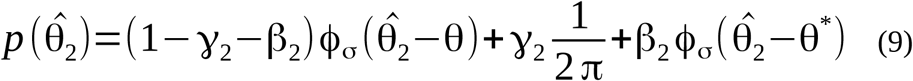

We were interested in the proportion of trials in which each of the two responses corresponded to the original memory item presented at the response location (**Equation 10-11**). This was obtained by excluding the proportion of swap errors *β* and the proportion of guesses *γ*.

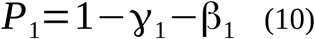

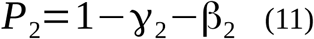

When participants are free to choose their response order in whole-report tasks, they show a decrease in recall accuracy as a function of response order, unlike participants whose response order was computer-determined (Adam et al., 2017). We thus introduced a parameter, asymptote *α*, to account for the difference in recall precision between the first and second response.

Since the response order does not necessarily reflect the order in which items are encoded, we applied the *α* parameter directly to the fitted probability of the second reported item being genuinely encoded, *P*_*2*_. The empirically obtained *P*_*2*_ includes only trials in which the second-report item was encoded and recalled, but not those in which it was encoded and then forgotten. The *α* parameter describes the proportion of trials in which the second-report item was encoded but consecutively forgotten. Thus, we can express the empirically obtained *P*_*2*_ as a proportion of the asymptotic recall, to account for post-encoding forgetting (**Equation 12**).

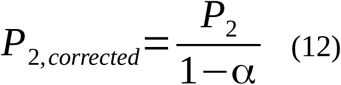

To prevent *P*_*2,proportional*_ from exceeding 1, we limited the upper bound of potential *α* values (**Equation 13**).

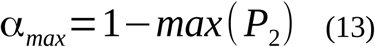

After obtaining *P*_*1*_ and *P*_*2,proportional*_ for each exposure duration, we could fit our encoding models (serial: **Equations 1-3 and 5-7**, parallel: **Equations 4 and 5-7**) to the resulting time series. In particular, we fitted the temporal profile of the probabilities of having encoded one or both of the items, by each moment in time during encoding. The three parameters, *α*, *κ*, and *t0* were estimated using least squares, to match the empirical time course of encoding probabilities to the model predictions. Note that *P*_*1*_ and *P*_*2,proportional*_ were obtained for every exposure duration, and that the sampling density was higher at lower exposure durations (six were under 120 ms, and three over 200 ms). Thus ordinary least-squares estimation will over-emphasise lower exposure durations. To overcome this, fitting was performed on interpolated datapoints resampled uniformly across exposure durations. Thus, *P*_1_ and *P*_*2, proportional*_ were linearly interpolating at every millisecond of exposure duration. These interpolated values were not counted towards the actual number of data points *n* that was used to compute the goodness of fit measures described next.

Goodness of fit was quantified with the *adjusted R*^*2*^ (**Equation 14**) and Bayesian information criterion (*BIC*, **Equation 20**). Differences in BIC are computed with respect to the best fitting model (which has the lowest BIC value), and interpreted according to Raftery’s guidelines (Raftery, 1995): ΔBIC values of 2-6 are considered positive evidence for the best fitting model being significantly better than the other models, 6-10 strong evidence, and values over 10 very strong.

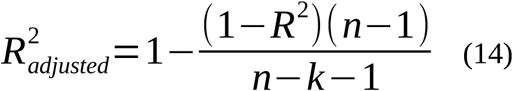

For which the un-adjusted *R*^*2*^ was computed via **Equation 15**.

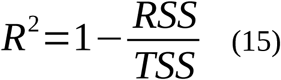

The residual sum of squares (*RSS*, **Equation 18**) and total sum of squares (*TSS*, **Equation 19**) were computed using the probabilities of only one (*P*_*one*_, **Equation 16**) or both items (*P*_*both*_, **Equation 17**) being in visual short-term memory. Specifically, the empirically derived probabilities *P*_*both*_ and *P*_*one*_ were compared with the estimated probabilities that both items (**Equation 5**) or only one item (**Equation 6**) were present in short-term memory. These estimated probabilities were based on either the serial (**Equations 1 and 3**) or the parallel encoding model (**Equation 4**).

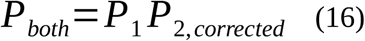

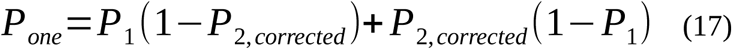

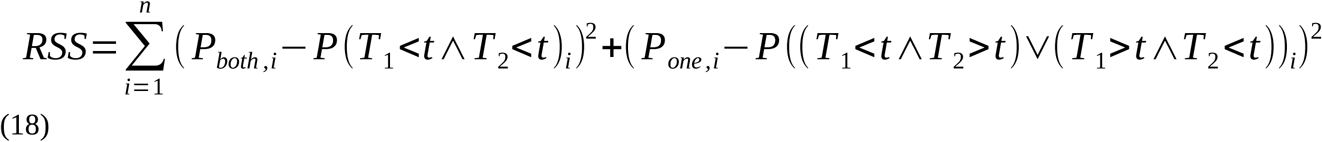

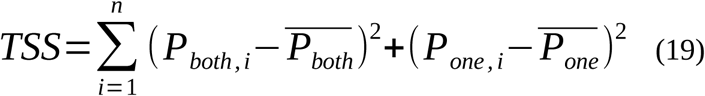

In a least-squares framework, the Bayesian information criterion (BIC) can be computed from the residual sum of squares (RSS).Our models were fitted on the same numbers of samples, and had the same number of free parameters, therefore, only the term including the RSS is relevant.

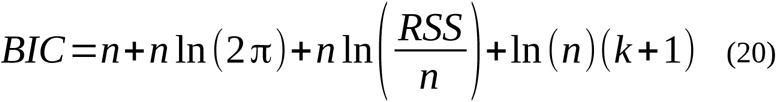

## Results

In our first experiment (N=11), two coloured disks were presented for 0, 10, 20, 30, 50, 90, 140, or 200 milliseconds, and masked directly after. After 500 milliseconds, participants could respond to both stimuli in the order of their choosing, by using the mouse cursor to click on a colour wheel around each placeholder. Error was computed as the unsigned circular distance between item colour and response. The probability of each item being in short-term memory was estimated by fitting a mixture model (Bays et al., 2009; Zhang & Luck, 2008) to the error distributions (see Methods). Error decreased as a function of exposure duration [F_greenhouse-Geisser_(2.47, 24.7)=102.89, p<0.001, η^2^=0.91], and was significantly lower for the first than the second response [F(1, 10)=79.00, p<0.001, η^2^=0.89], but only for exposure durations of 50 milliseconds and higher [F(4, 40)=11.34, p<0.001, η^2^=0.53].

Because all trials had two items, participants could pre-allocate their attention to one location. In addition, our first study lacked an independent estimate for the first item to be encoded. Therefore, we performed a second experiment (N=11) in which we introduced a single item condition in which a single coloured disk could be presented on either stimulus position. We also extended the range of exposure durations to 0, 10, 20, 40, 70, 120, 200, 360, and 580 milliseconds. As expected, error decreased as a function of exposure duration [F(8, 80)=178.24, p<0.001, η^2^=0.95], and was significantly lower for the single and first than the second response [F(2, 20)=39.78, p<0.001, η^2^=0.80], but only for exposure durations of 40 milliseconds and higher [F(16, 160)=19.24, p<0.001, η^2^=0.66]. Importantly, no difference in error existed between the single-item condition and the first-reported item in the two-item condition [*p*_*Holm*_=0.156], suggesting that the encoding of the second item did not slow down that of the first.

To determine whether our findings generalised to other kinds of information, our third experiment (N=16) used memory for visual orientation instead of colour. The design was identical to the second, with the exception that items were greyscale sinusoidal gratings with a clear orientation. Again, error decreased as a function of exposure duration [F(8, 120)=147.77, p<0.001, η^2^=0.91], and was significantly lower for the single and first than the second response [F(2, 30)=68.59, p<0.001, η^2^=0.82], but only for exposure durations of 40 milliseconds and higher [F(16, 240)=17.89, p<0.001, η^2^=0.54].

Previous research has demonstrated load-dependent activity in left parietal cortex only for items in the right hemifield, while activity in right parietal cortex reflected items across the entire visual field (Sheremata, Bettencourt, & Somers, 2010), suggesting a degree of independence between cerebral hemispheres. It is therefore possible that parallel encoding in our experiments depends on the items being presented in different hemifields. To rule out this explanation for our results, we conducted a fourth experiment (N=22), in which two stimuli were shown either unilaterally or bilaterally to investigate multi-item encoding within the same hemisphere. As before, error decreased as a function of exposure duration [F_Greenhouse-Geisser_(2.59, 54.32)=332.88, p<0.001, η^2^=0.94], and was significantly lower for the first than the second response [F(1, 21)=72.06, p<0.001, η^2^=0.77], but only for the exposure durations of 70 milliseconds and higher [F_Greenhouse-Geisser_(4.65, 97.72)=19.05, p<0.001, η^2^=0.48]. In keeping with research showing a degree of hemispheric independence, errors were marginally but significantly lower for bilateral than unilateral presentation [F(1, 21)=7.87, p=0.011, η^2^=0.27], but only at higher exposure durations [F(8, 168)=2.67, p=0.009, η^2^=0.11] (due to the lack of a difference at lower exposure durations).

In all experiments, encoding probabilities were better fitted by the parallel than the serial model (**Table 1**). Collapsed across bilateral presentation conditions in experiments with the same exposure durations (N=49), the parallel model fitted better than the serial model (**Figure 2**). Parameter estimates aligned well with previous research (Bundesen, 1990; Vogel et al., 2006) for both models: parallel *κ*=19.94 Hz, *t0*=5.69 ms, *α*=0.29; serial *κ*=14.73 Hz, *t0*=0.09 ms, *α*=0.29.

**Table 1.** Goodness-of-fit measures across experiments and conditions for the serial and parallel models fitted to joint encoding probabilities. BIC differences are computed with respect to the best-fitting model, which was the parallel model in all experiments.

| | Encoding time (ms) | Serial $R^2_{\text{adjusted}}$ | Parallel $R^2_{\text{adjusted}}$ | $\Delta\text{BIC}$ |
| --- | --- | --- | --- | --- |
| **Experiment 1** (bilateral, colour) | 0-200 | 0.84 | 0.88 | 1.95 |
| **Experiment 2** (bilateral, colour) | 0-580 | 0.89 | 0.89 | 0.95 |
| **Experiment 3** (bilateral, orientation) | 0-580 | 0.89 | 0.93 | 3.69 |
| **Experiment 4** (unilateral, colour) | 0-580 | 0.92 | 0.94 | 2.46 |
| **Experiment 4** (bilateral, colour) | 0-580 | 0.89 | 0.92 | 2.62 |
| **All bilateral conditions** (sum) |  |  |  | **9.21** |

**Figure 2.**
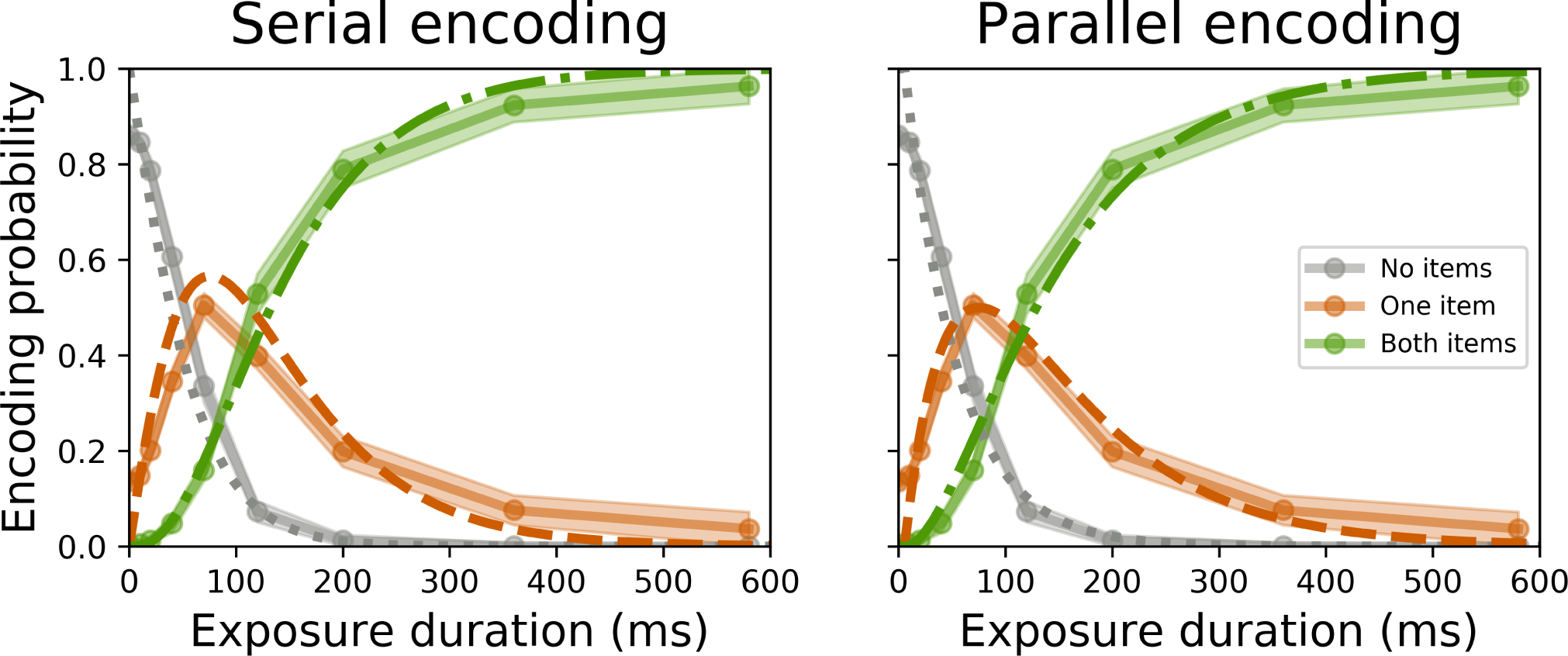
The plots show averages (solid lines) and standard error (shading) estimated from empirical data of the encoding probabilities of no (grey), only one (orange), or both items (green). The best fitting solutions of the serial (left) and parallel (right) models are drawn in dashed and dotted lines, and show that the parallel model fitted the data best. These data were collected in the bilateral presentation conditions of Experiments 2-4, from a total of 49 participants.

Although for each experiment the ΔBIC values do not seem particularly high, taken together the four experiments provide strong evidence for the parallel model. It should be noted that both models fitted very well, which limits the practical range of BIC differences. Crucially, even in theory the difference between the two models should only be obvious at the cirtical exposure durations when it is likely that only one of the two items has been encoded. This greatly limits the number of data points that can contribute to BIC differences. Moreover, different subjects have different times at which this critical peak occurs. Despite these limitations, all BIC differences were consistently in the direction of the parallel model, and nearly all were in a range that is considered positive evidence (Raftery, 1995).

## Discussion

We conducted four whole-report experiments in which we briefly presented items, and asked participants to recall and reproduce all of them. Recall accuracy increased as a function of exposure duration, replicating earlier work (Bays et al., 2011; Bundesen, 1990; Vogel et al., 2006). In addition, recall accuracy was slightly better when items were presented bilaterally rather than unilaterally. Crucially, we found that when two items were presented, the probability of only one of them being encoded into visual short-term memory did not peak as sharply as one would expect under a serial encoding model. Instead, our results were better fitted by a parallel encoding model, suggesting that information from several items can be encoded independently and at the same time.

One recent study attempted to compare the serial and a parallel encoding hypotheses by employing serial and simultaneous stimulus presentation, and came to the opposite conclusion (Liu & Becker, 2013). Unfortunately, the experimental design of that study failed to account for rate-limitations of encoding: It included imbalanced exposure durations for serial (150 ms per stimulus) and simultaneous presentation (150 ms for both stimuli). In addition, the employed models were labelled ‘serial’ and ‘parallel’, but did not incorporate encoding time. Notably, the ‘parallel’ model lacked the inter-item independence that is characteristic of parallel encoding: a guessing rate parameter was fixed between items. In our opinion, the experiments and models in the current study better capture and characterise the encoding process, as they investigate encoding probability as a function of encoding time, and crucially allow for items to be independently encoded.

Although we provide evidence for parallel encoding of information into short-term memory, whether humans are able to encode information in a serial fashion remains an open question. The methods and models presented here could be used in future studies to address this, for example by incentivising the precise recall of a particular item within a whole-report trial by reward. The reward-associated item could be encoded before all others, or encoded at the same time but at a higher rate. With minor adjustments, our models would be able to differentiate between these hypotheses. Specifically, the serial model should allow for reward-dependent encoding order, and the parallel model for reward-informed bias in encoding rates.

## Conclusion

In conclusion, our results provide evidence that at least two items can be encoded into short-term memory in parallel. This suggests a reformation of the popular feature-integration theory of attention (Treisman & Gelade, 1980) and short-term memory (Wheeler & Treisman, 2002) could be in order, specifically where it postulates serial binding and encoding. Additional implications pertain to two-stage cognitive models that incorporate feature-integration theory. These generally propose an initial pre-attentive parallel perceptual processing that is followed by a stage in which attention is serially deployed to bind information within each independent item (Cave & Wolfe, 1990; Duncan, 1984; Hoffman, 1979; Treisman, 1982). Contrary to these models, others have demonstrated that attention can be deployed in parallel to non-contiguous locations (Hahn & Kramer, 1998; Kramer & Hahn, 1995; McMains & Somers, 2004; Müller et al., 2003), and the current study suggests this is also true for short-term memory encoding.

We propose that attentional resources can be distributed (quantised or fluidly) among several items to form independent encoding channels that consolidate information into the same central short-term memory store. Alternatively, one channel could (rapidly) alternate between several stimuli, partially encoding each. This speaks against computational models that assume serial complete encoding, and illustrates the biological plausibility of models that allow for simultaneous encoding of several items (e.g. (Matthey et al., 2015)).

## Acknowledgements

ESD was supported through a European Union FP7 Marie Curie ITN grant (606901). SGM is supported by an MRC Fellowship MR/P00878X/1. MH is supported by the Wellcome Trust and the NIHR Oxford Biomedical Research Centre. We thank Rachel King and Chiron Oderkerk for fruitful discussions on the subject of short-term memory encoding, and Gemma Lowcock for helping with data collection for experiment 4.

## Author contributions

E.S.D. and M.H. initiated the project, conceived the ideas, and designed the experiments. E.S.D. collected the data. E.S.D. and S.G.M. conceived the analysis. E.S.D. analysed the data, and wrote the manuscript with contributions from all authors.

